# Ensembles from ordered and disordered proteins reveal similar structural constraints during evolution

**DOI:** 10.1101/468801

**Authors:** Julia Marchetti, Alexander Miguel Monzon, Silvio C.E. Tosatto, Gustavo Parisi, María Silvina Fornasari

**Affiliations:** Departamento de Ciencia y Tecnología, CONICET, Universidad Nacional de Quilmes, Roque Sáenz Peña 352, B1876BXD Bernal, Provincia de Buenos Aires, Argentina; Department of Biomedical Sciences, University of Padua, Padua, Italy

**Author notes:** Corresponding author: Gustavo Parisi.

**Keywords:** protein evolution, protein ensemble, conformational diversity, disordered proteins

## Abstract

Inter-residue contacts determine the structural properties for each conformer in the ensembles describing the native state of proteins. Structural constraints during evolution could then provide biologically relevant information about the conformational ensembles and their relationship with protein function. Here, we studied the proportion of sites evolving under structural constraints in two very different types of ensembles, those coming from ordered or disordered proteins. Using a structurally constrained model of protein evolution we found that both types of ensembles show comparable, near 40%, number of positions evolving under structural constraints. Among these sites, ~68% are in disordered regions and ~57% of them show long-range inter-residue contacts. Also, we found that disordered ensembles are redundant in reference to their structurally constrained evolutionary information and could be described on average with ~11 conformers. Despite the different complexity of the studied ensembles and proteins, the similar constraints reveal a comparable level of selective pressure to maintain their biological functions. These results highlight the importance of the evolutionary information to recover meaningful biological information to further characterize conformational ensembles.

## Introduction

The protein native state is described by a collection of the different structures which a given sequence could adopt. This collection is also called a conformational ensemble and is an essential concept to understand protein biology ^1,2^. The existence of conformational ensembles is known since the crystallization of hemoglobin with its two conformational states T and R (deoxy and oxygenated forms) in the early 1960. The growth of Protein Data Bank (PDB) redundancy, refinement and development of techniques such as NMR, SAXS and single molecule spectroscopy over the last years have allowed the experimental characterization of a large number of protein ensembles ^2,3^. Structural differences between conformers could result from the relative movements of large domains as rigid bodies ^4^, secondary and tertiary element rearrangements ^5^, and loop movements ^6^. Apparently, most globular proteins have very few conformers describing their native state to achieve their functions. Proteins with little flexibility at the backbone level, called rigids, have only one conformer in their ensembles ^7^ like the cellulase from *C. cellulolyticum* ^8^. Hemoglobin, as mentioned previously, is the paradigm for proteins with two conformers ^9^, while the dimeric catabolite activator protein ^10^ and the human glucokinase have three ^11^. Complex proteins composed of several different chains, like mitochondrial ATP synthase could have at least seven conformers ^12^. As protein flexibility increases, the number of conformers in the ensemble increases as well, giving rise to very complex ensembles as in the case of intrinsically disordered proteins (IDPs) or regions (IDRs). IDPs are characterized by the lack of tertiary structure under physiological conditions ^13,14^. IDP ensembles are composed by a large number of interconverting conformers given their low free-energy barriers among them ^15^. Far from being random polymers or random-coiled ensembles, it is becoming evident that IDP ensembles are not fully disordered, showing transient short and long-range structural organization ^16^. Order-disorder transitions are frequently observed in IDPs or IDRs, sometimes associated with ligand binding ^17^ but in other cases just reflecting the heterogeneous composition of the ensembles ^7,18^.

Here, we studied the level of structural constraints in IDPs ensembles compared with those found in globular proteins. Structural constraints could be studied using direct methods such as the measurements of contacts between residues in a given structure or conformer and some derived parameters such as the contact density (mean number of residue-residue contacts per residue) or their interaction networks ^19^. However, inter-residue contacts could be artifacts or simply be irrelevant in very complex ensembles such as those found in IDPs, making it difficult to detect biologically relevant conformers ^20^. For these reasons, in this work we evaluated the amount of structural constraints using an evolutionary approach. It is a well-established concept that conservation of protein structures during evolution constrains sequence divergence modulating in this way the amino acid substitution pattern of certain positions ^21,22^. These structural constraints are evidenced in sequence alignments as differentially conserved positions, showing a given physicochemical bias or subject to coevolutionary processes due to their relative importance to maintain protein fold and dynamics (i.e. conservation of given interactions to increase stability, sustain protein movements). This structurally constrained substitution pattern has been exploited to improve models of molecular evolution ^23–25^, explain rate heterogeneity ^26^, make functional predictions ^27^, compare the substitution process in ordered and disordered proteins ^28^ and in the inference of given tertiary folds ^29^ to mention just a few examples of their many applications. Furthermore, evolutionary information could be used to predict native contacts and structural models of globular domains ^30–32^. More recently these methods were adapted to successfully predict globular states in disordered proteins and to show the evolutionary constraints in protein interfaces between disordered and ordered proteins again showing the importance of structurally constrained information during evolution ^33,34^. Substitution patterns observed in sequence alignments can be described by evolutionary models ^35^. Alternative models, making different assumptions about the amino acid substitution pattern, can be compared using maximum likelihood estimations to decide which assumptions better describe the evolutionary process in a given family. In particular, in this work a model of protein evolution using protein structure to derive a structurally-constrained site-specific substitution pattern was used ^24^. As this models is structure-specific each protein conformation represents different evolutionary models. Using maximum likelihood estimations, we then compared how the structurally-constrained substitution pattern outperforms models of evolution lacking structural information (e.g. JTT ^36^, Dayhoff ^37^, WAG ^38^) in its ability to explain the observed site-specific substitution pattern in a set of homologous proteins for each studied protein. Interestingly, considering all conformers in the ensembles of globular and IDP proteins, we found that the number of structurally constrained positions are similar for both kinds of proteins.

## Results

### Description of the datasets

In the last years, an emerging picture evidences that increasing structural differences between conformers, connected by very different dynamical behaviours, produces a continuum in protein space ^39^. One extreme feature of this continuum is the presence of rigids proteins with almost no backbone differences among their conformers and just displaying only conformational diversity at the residue level ^7^. Increasing conformational diversity at the backbone level could evidence the presence of disorder, where the appearance of short-time dynamical behaviour allows the sampling of a large conformational space ^40^. Figure 1 shows different types of ensembles as protein conformational diversity increases. In one extreme of the distribution (left-side panel in Figure 1) typical globular or ordered proteins are shown. These proteins generally show large proportions of secondary structure where their spatial arrangement defines a single tertiary structure and hydrophobic core. The higher density of inter-residue interactions of this core constrains evolutionary rates when compared to exposed residues ^41^ and also contains enough information to define a global tertiary arrangement ^42^. As mentioned before, ordered proteins could also contain different conformers to achieve their biological functions (Figure 1, middle-panel), giving place to additional restrictions in the protein substitution pattern ^43^. Middle-panel examples of Figure 1 could also display proteins with ordered or globular regions as well as very flexible regions showing different dynamical behaviour and possibly originating disordered regions of different lengths. Right panel in Figure 1, shows a typical ensemble of IDPs showing a collection of conformers determined by NMR. These ensembles show highly flexible chains and eventually small and transient segments of secondary or tertiary structure ^44^. Consequently, IDPs have a large degree of conformational entropy that can be limited by inter-residue interactions originating a complex mixture of conformers in the ensemble ^15,20^. As described in methods, two hand-curated datasets were analysed. The ordered dataset composed of 227 proteins with known crystallographic structure containing non missing residues and a disordered dataset containing 93 NMR ensembles of different proteins. Disorder has been estimated in both datasets using ESpritz and Mobi 2.0 for the disordered and ordered datasets respectively (See Methods). As is it shown in Figure 2, ordered proteins show a low predicted content of disordered residues while the disordered dataset shows a distribution of disordered residues. The median of these distribution is 58% of disordered positions (minimum 40% and up to 98%). It is then expected that the disordered dataset contains small globular regions and more than the half of the protein in a disordered state. Sequence alignments for each protein in each dataset were extracted from HSSP database (see Methods) and to avoid high occurrence of indels, sequences above 30% identity with the protein with known structure were only considered. Additional information about protein alignments could be found in Figure S1.

**Figure 1:**
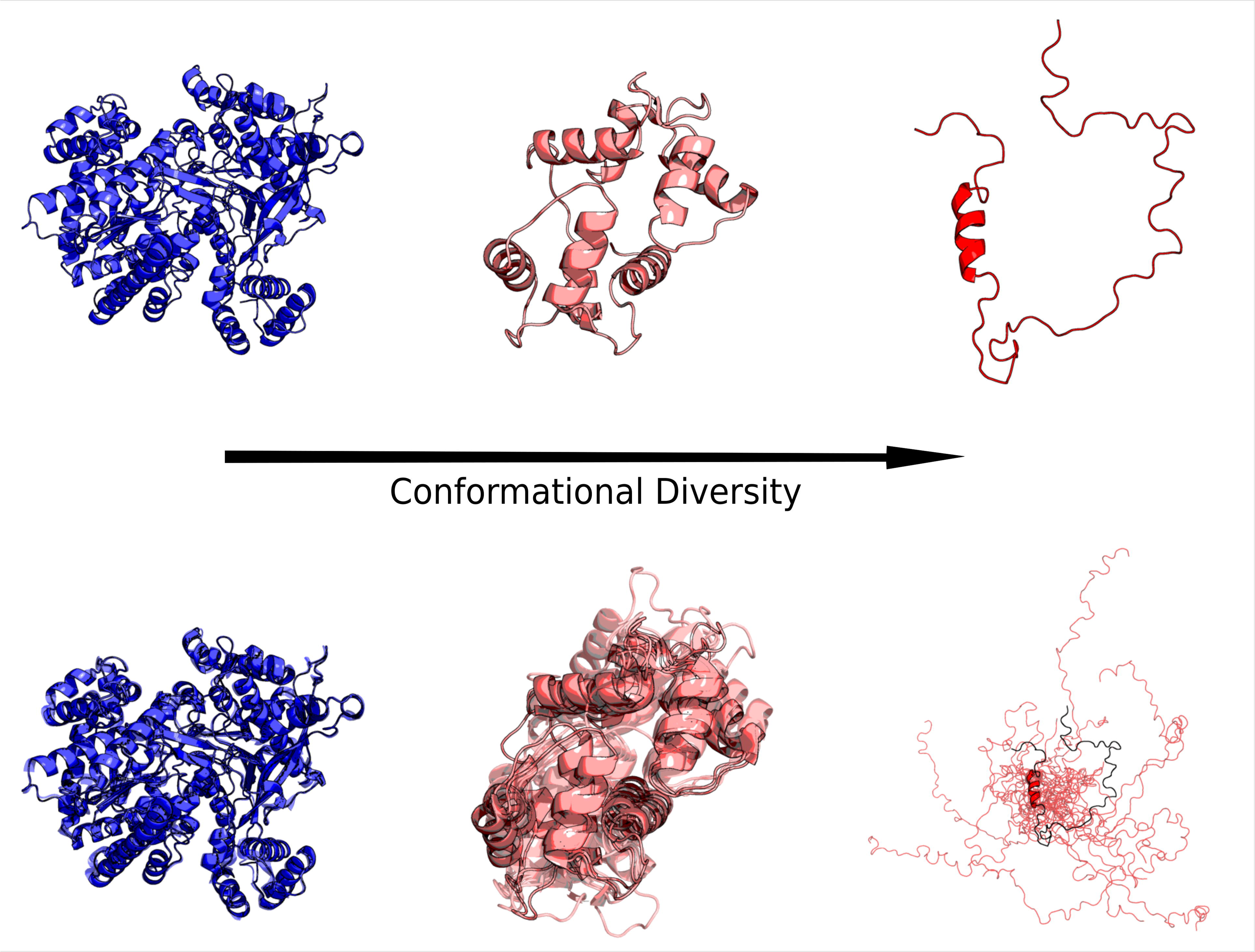
Different protein ensembles as a function of flexibility increases. Top panel shows a given conformer while the bottom panel shows all the available conformers in the ensemble. Left, maltodextrin phosphorylase, (PDB codes = 1AHP_A, 1AHP_B, 1L5V_B) shows as a rigid protein with 6.53% disordered and it is taken as a representative of ordered proteins. Calmodulin (PDB codes = 2FOT_A, 1LIN_A,1NIW_E, 3G43_A, 2BE6_A, 1CDL_A, 3GP2_A, 4L79_B, 1CLL_A) shows 10.64% of disorder. Thylakoid Soluble Phosphoprotein, (PDB ID = 2FFT_A) is a typical IDP ensemble with 100 percent of estimated disorder. The percentages of disorder were estimated with ESpritz.

**Figure 2:**
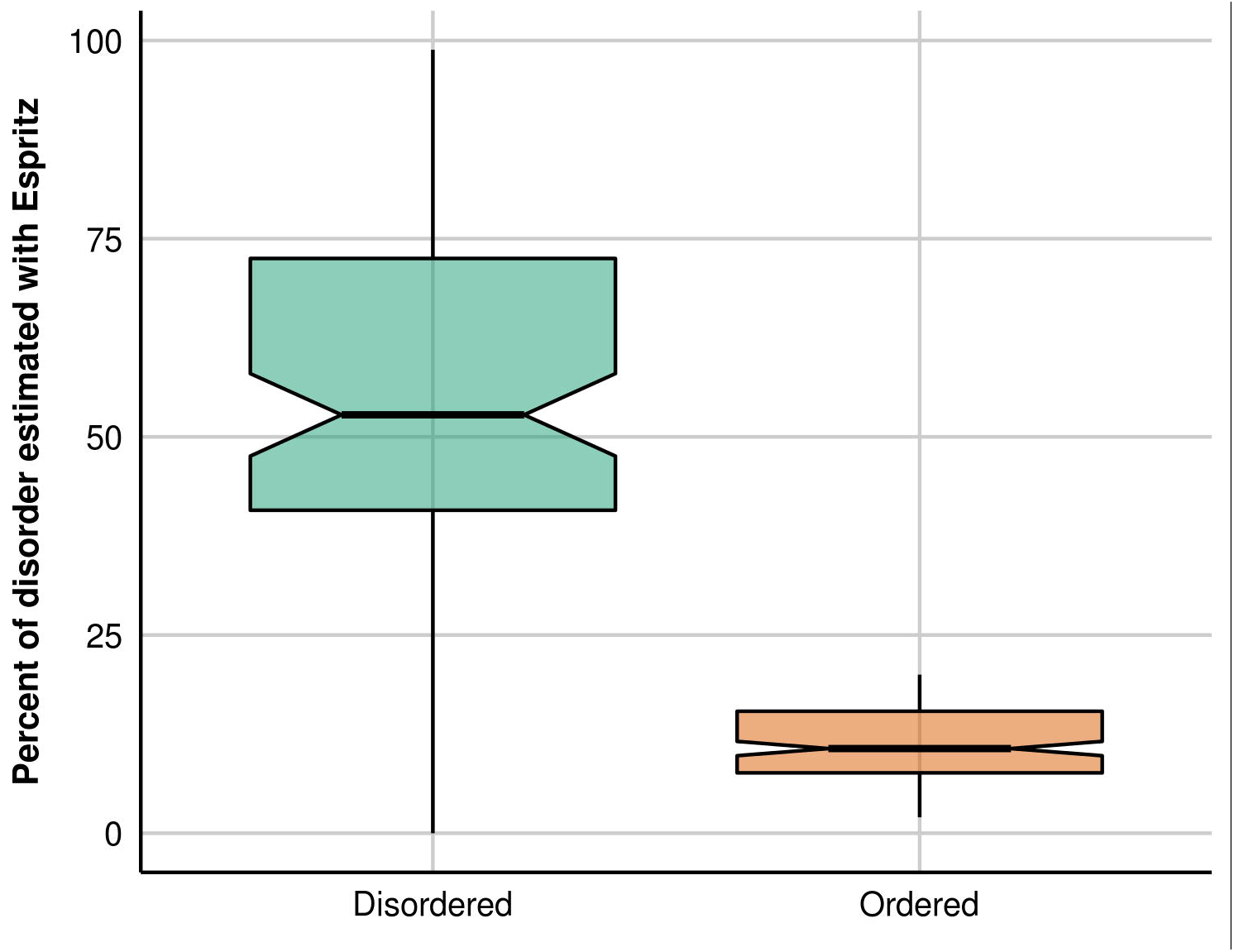
Estimation of disorder content using NMR-ESpritz in the disordered set and ESpritz in the ordered set. It is shown that the ordered set has a low proportion of disorder well below the reported error in the estimation ^58^.

### Physical contacts versus structural constraints during evolution

To assess the structural constraints in ordered and disordered ensembles, we quantified the inter-residue interactions accumulating the contact information for each site and protein through all the available conformers in each corresponding ensemble (Figure S2, panel A). Accumulation is a reasonable idea sustained by the particular contributions each conformer makes to the biological function ^2^. As a result, we obtained that the great majority of residues are involved in inter-residues contacts as it is shown in Figure 3a. Permanent secondary and tertiary contacts in ordered proteins define their levels of structural constraints while the contribution of transient contacts along the entire ensemble of IDPs produces almost the same amount of accumulated inter-residues contacts (3rd quartile is 100% and 97% for IDPs and ordered sets respectively). According to this result the vast majority of positions in IDPs are constrained by structural restrictions as well as those for ordered proteins. However, it is well established that IDPs have a different pattern of amino acid substitutions when compared with ordered proteins. IDPs show also a highly conserved composition of amino acids ^45^ instead of the well defined site-specific substitution pattern observed in ordered proteins ^46^. Mostly, IDPs and IDRs show higher evolutionary rates as well as higher rates of insertions and deletions compared with their ordered counterpart ^44^ ^<u>1347</u>^. To elucidate the influence of such high levels of structural constraints (Figure 3a), we turned to study the substitution pattern observed in the homologous family of each protein in both datasets. Using maximum likelihood comparisons (See Figure S2, panel B), we assessed if the observed substitution pattern is better explained by a evolutionary model containing structural information (like SCPE, see Methods) or by other models not containing this information (JTT, Dayhoff and WAG models, see Methods). When SCPE site-specific substitution matrices outperform each of the three other models, it is inferred that the site evolves under structural constraints. Considering the different nature of ordered and disordered ensembles, unexpectedly, we found that the percentages of structurally constrained sites (SC) is almost the same in both types of ensembles (41.6% and 40.5% for disordered and ordered datasets; Figure 3b) and much lower than estimations made using the accumulated account of inter-residue contacts. Interestingly, the individual conformers show slightly less percentages of SC sites (Figure 3c) showing 32.1% and 36.1% in average for the disordered and ordered datasets.

**Figure 3:**
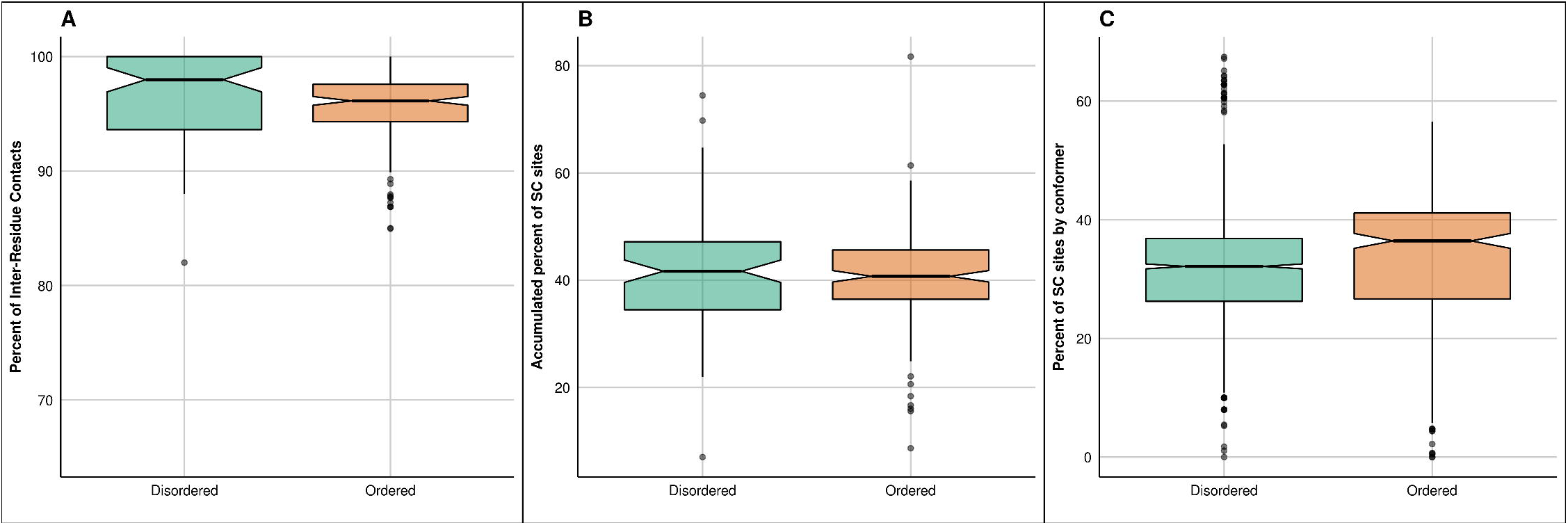
(A) Percentage of inter-residue contacts for the disordered and ordered datasets (average median of 96.1%). (B) Distribution of the accumulated number of structurally constrained sites for both datasets showing 41.6 and 40.5% of the positions. The distributions are statistically similar using a Kolmogorov-Smirnov test with p-value = 0.39 and Mann-Whitney-Wilcoxon test with p-value = 0.45. (C) Distribution of structurally constrained sites per conformer per protein showing a median of 32.1% and 36.1% of their sites constrained.

### Structurally constrained sites

SC sites are then sites that at least have one physical inter-residue contact in at least one conformer but also, and more importantly, modulates sequence divergence in that specific position. To further investigate these structural constraints we studied the distribution of SC sites. We found that ~68% of the SCs are located in the disordered regions of the proteins belonging to the disordered dataset (Figure 4). As we mentioned before, disordered proteins could have permanent or transient globular regions that could increase the structural constraints of the protein as a whole. However, the number of SC sites in the globular or ordered regions of the disordered proteins is ~32%. This results indicate that globular regions of disordered proteins are less constrained that the corresponding in the ordered dataset (see Figure 3b). Also, following our definition of inter-residue contacts (see Methods), all estimated contacts are tertiary and in ~57% the SCs are classified as long-range inter-residue contacts (see Figure 5). This finding can explain how SC sites could appear in disordered regions. As we can see in Figure 6 disordered proteins could have large conformational diversity. However, among the representatives conformers of the ensembles we can find some of them collapsing over the globular part of the protein or just adopting close conformations increasing in this way the number of contacts per site. As it is shown in Figure 7, 51% of the positions have contacts that are present in the 100% of the conformers of the ensemble. However, there is still a tail in the distribution showing that single conformers could have SC sites, in other words, single conformers could have inter-residue contacts that modulate the substitution pattern of those positions.

**Figure 4:**
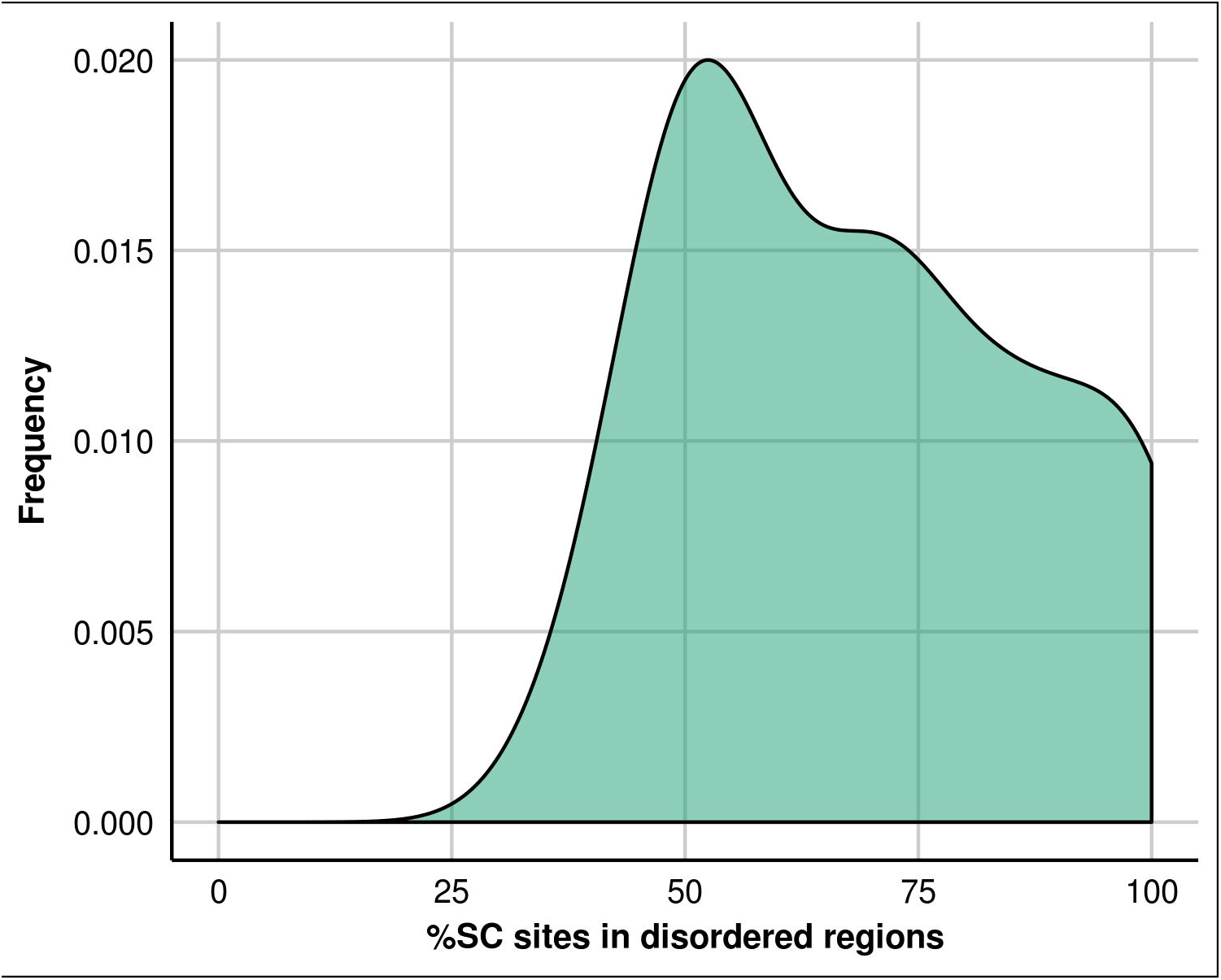
Distribution of the accumulated number of structurally constrained sites along all the ensemble. On average 68.3% of the SC sites are within predicted disordered regions.

**Figure 5:**
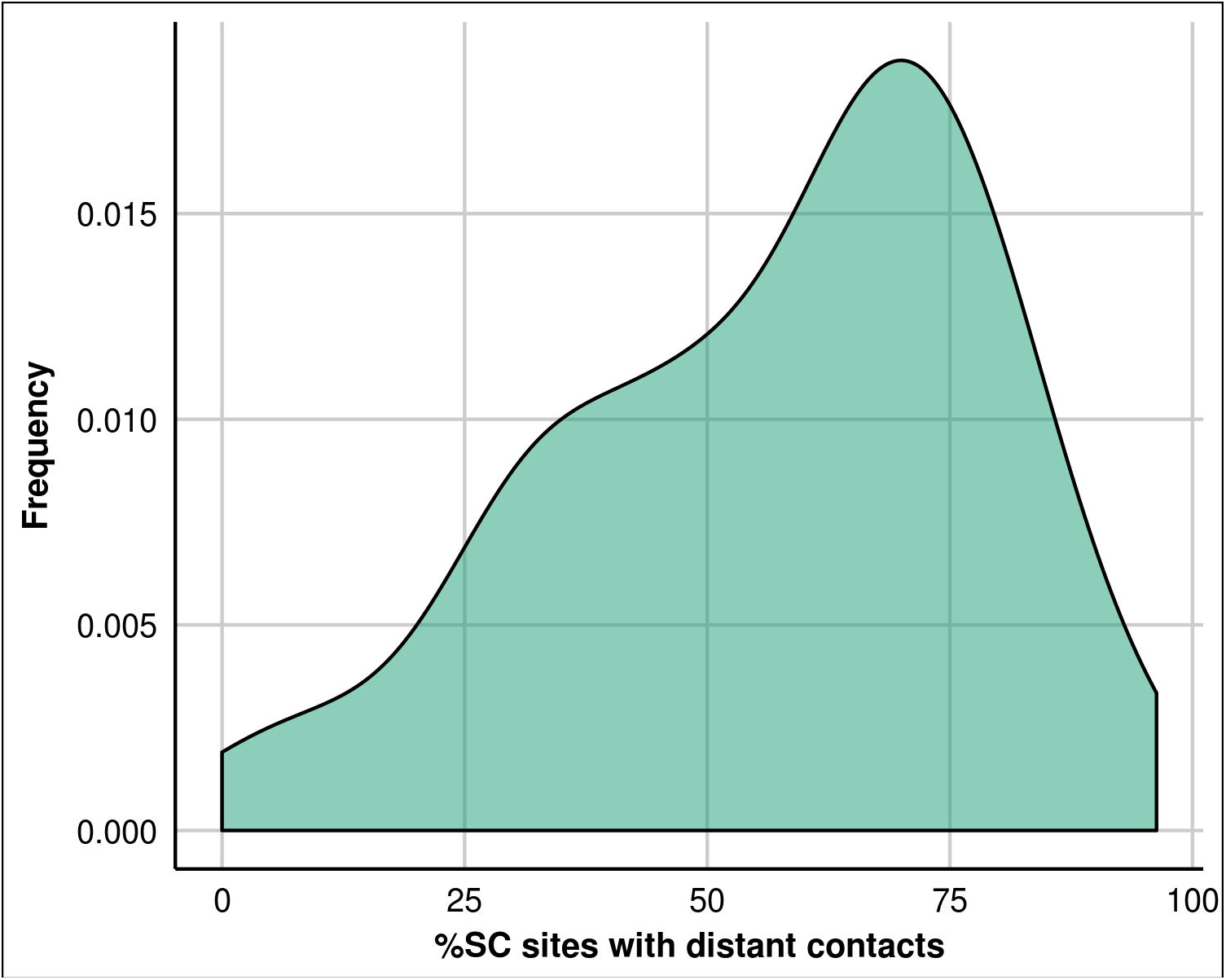
Distribution of the accumulated number of structurally constrained sites along all the ensemble, with long-distance contacts (at least 5 residues away). In average 56.8% of the SC sites have long-range inter-residue contacts.

**Figure 6:**
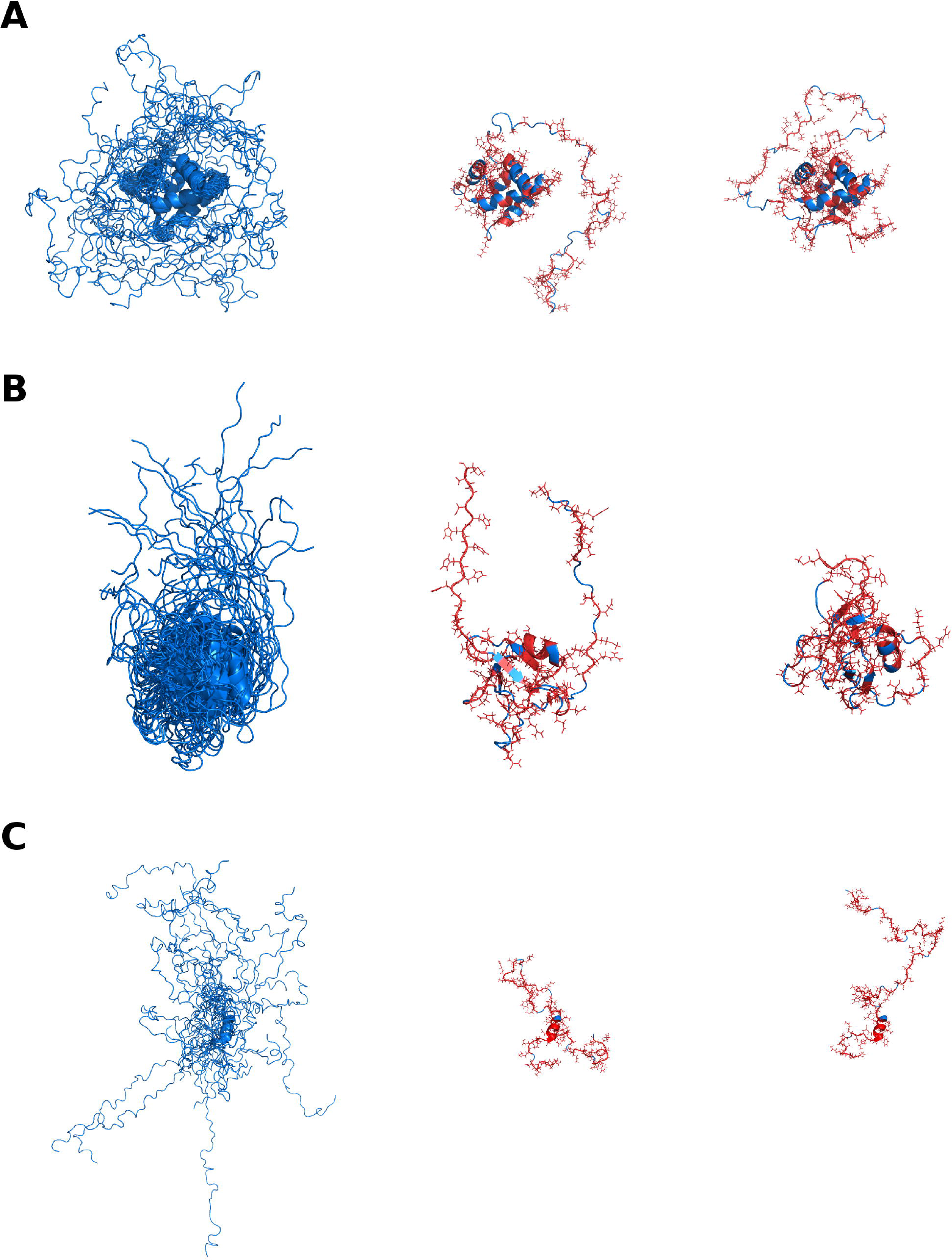
Examples showing SC sites distribution in different conformers. The three panels (top, middle and bottom) contain disordered proteins showing in the left the available ensemble, while in the middle and in the right we show different conformers. Proteins are shown in cartoon representation in blue while SC sites are shown in red sticks. 2JRF_A, 2ADZ_A and 5MRG_A are the corresponding PDB codes for the top, middle and bottom panels.

**Figure 7:**
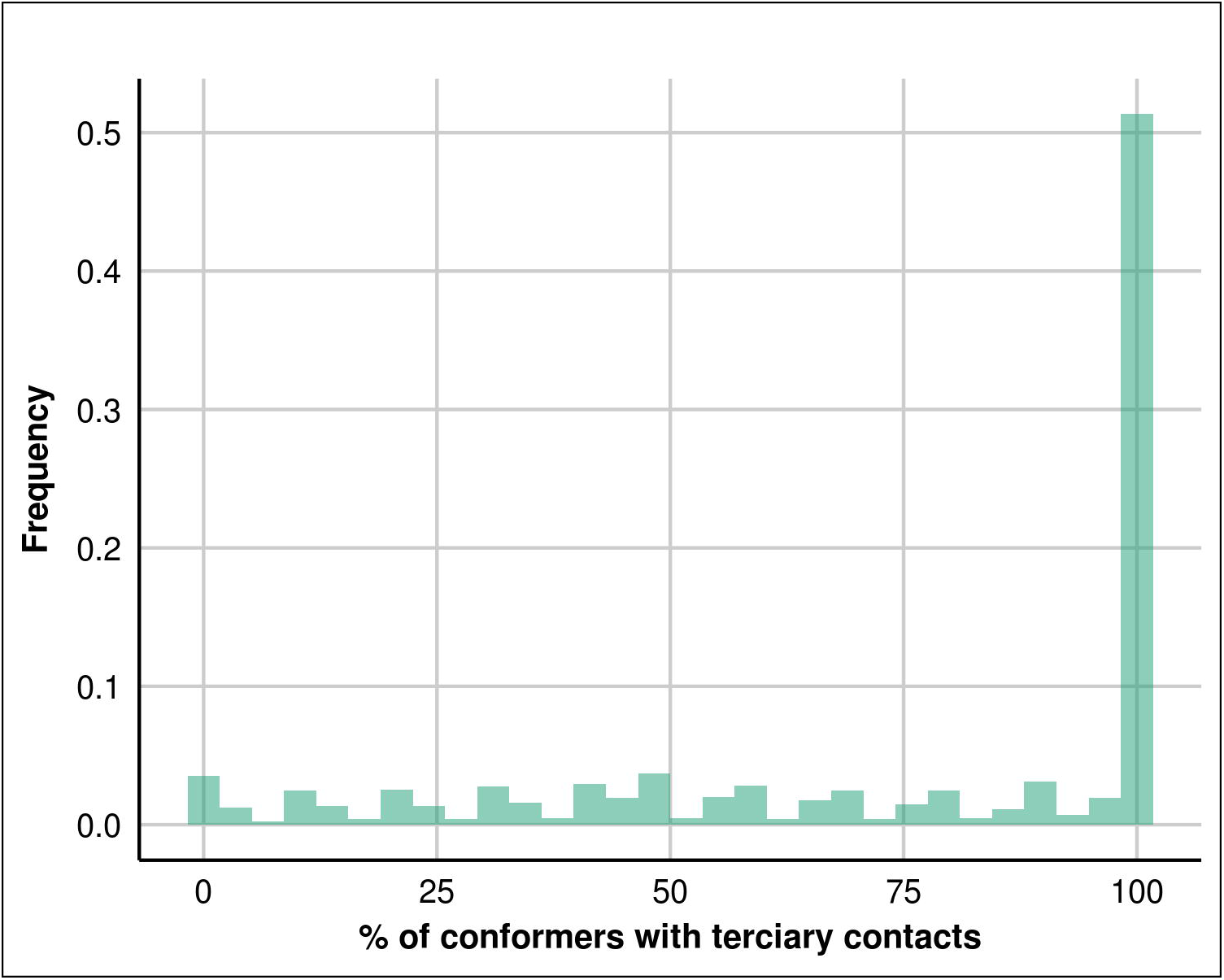
Approximately ~51% of SC sites present contacts in 100% of the conformers and only ~3% of SC sites present contacts in 50% of the conformers.

### Ensemble redundancy

How many conformers are required to fully describes evolutionary structural constraints contained in sequence alignments? When we calculated the minimum number of conformers per ensemble to reach the accumulated SC percentage per protein, we found that on average ~11 conformers are required for the proteins in the disordered dataset (see Figure 8) while in ordered it is ~1.5. The value for the ordered dataset is consistent with the available experimental evidence. Most ordered proteins show low conformational diversity, and then are called “rigid” ^7^, or could show very few conformers, mostly two referring to the bound and unbound forms of the protein^<u>48,4950</u>^. Due to the complexity of disordered ensembles, the number of conformers is difficult if not impossible to estimate. However, our measure of the number of conformers required to explain the evolutionary structurally constrained information in sequence alignments could offer a proxy to the number of conformers. Since the average of conformers in the NMR ensembles in our dataset is ~20, our results indicate that are mostly redundant.

**Figure 8:**
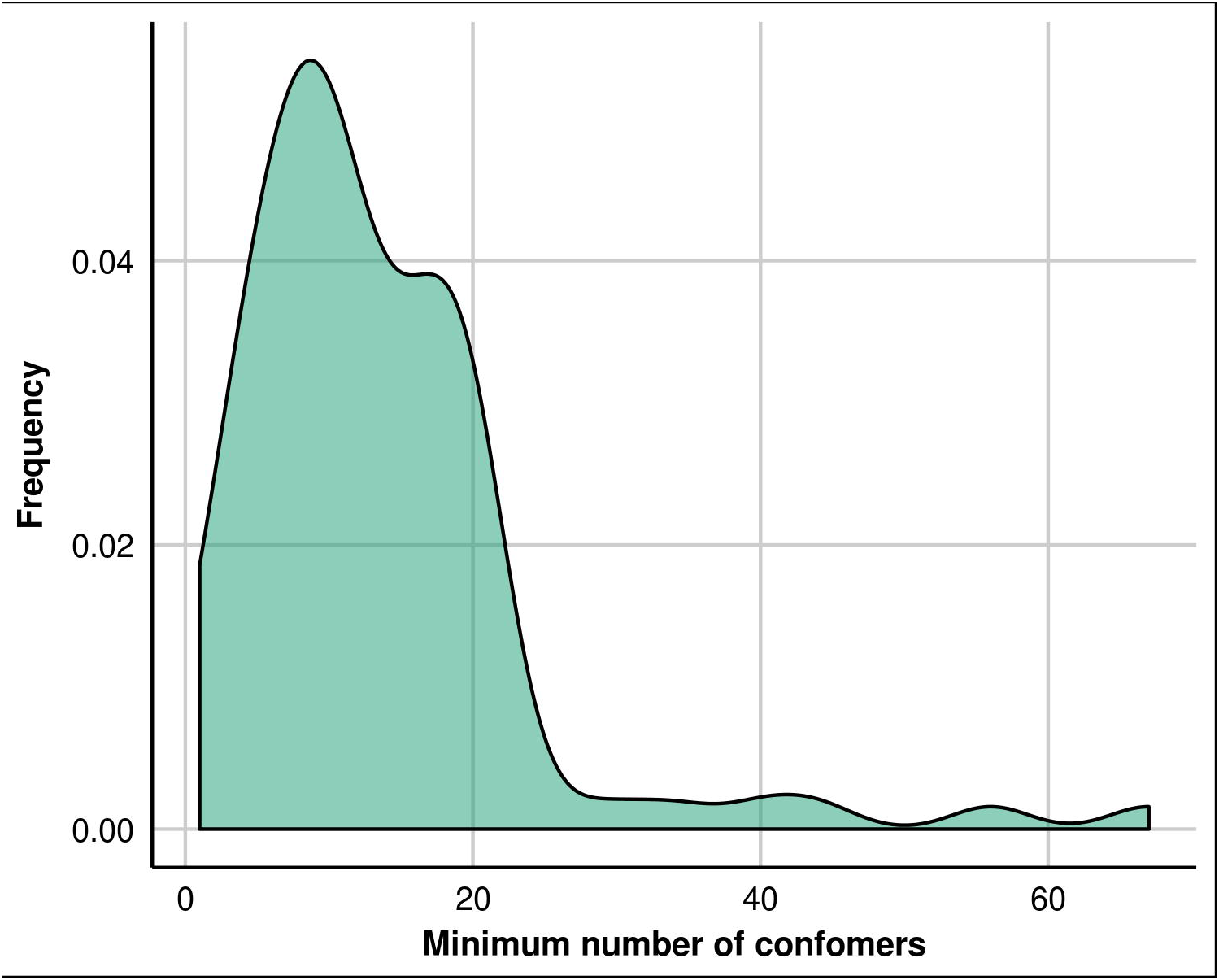
Distribution of the minimum number conformers to reach the accumulated percentage of structurally constrained (SC) sites per protein for the 93 disordered proteins corresponding to the set obtained with Mobi 2.0 and ESpritz (NMR). Minimum = 1, average ~11 and maximum ~64.

## Discussion

Two main findings emerge from the present work. First, the number of positions having inter-residue contacts accumulated along all available conformers in each ensemble approaches almost 100% of the positions (Figure 3a). However, as we have shown, the average percentage of positions evolving under structural constraints is much lower ~40% (Figure 3b). Part of this reduction is expected, given that not all intramolecular non covalent contacts could be equally relevant, for example, in structure stabilization ^51^. Additionally, the reduction could be also attributed to the lack of protein-specific information contained in sequence alignments. This effect operates over SCPE substitution matrices which are site and conformer specific but are evaluated using sequence alignments from corresponding homologous families. Thus, evolutionary information contained in those alignments reflects constraints of several sorts, such as structural divergence ^41^ or dynamical adaptations ^52,53^ which could certainly modify the contact pattern in the homologous proteins. It is then expected that this ~40% of structurally constrained sites on average obtained for both ensembles does not capture subtle inter-residue contacts originated in functional adaptations for individual proteins. In line with this observation, it has been recently shown that the use of sequence alignments recovers the most conserved pattern of inter-residues contacts when co-evolutionary and evolutionary coupling methods are used ^53^. The other important result is related with the similar structural constraints on sequence divergence in ordered and disordered proteins (Figure 3b). Our results suggest that individual contributions of each conformer in the disordered ensemble are required to sustain biological function as is well established for ordered proteins, and more recently suggested for disordered ones ^2,13,47^. These small contributions from each disordered conformer give overall the same proportion of structural constraints as found in ordered proteins, possibly with different weights according to their biological role.

Interestingly, the number of conformers in the IDPs ensembles to reach the corresponding level of global constraints per protein is ~11 (Figure 8). This means that IDP ensembles are redundant in terms of conformations and that possibly the number of relevant conformers in IDP ensembles would not be so large as expected due to their higher flexibility. These results are in agreement with the idea that different members of the ensemble could be directly involved in protein function but also they could be important as a local minimum representatives in the interconversion of biologically relevant conformations ^54^.

Our results highlight the importance of the evolutionary analysis in the discrimination of inter-residue contacts to detect meaningful biological information as well as the estimation of the number of conformers and structural constraints in such complex ensembles as those belonging to IDPs.

## Materials and Methods

### Dataset collection

Globular or ordered protein ensembles were obtained from the CoDNas database ^55^. Considering the presence of missing residues as a primary indicator of IDRs in proteins ^56^, we selected 227 proteins having no missing residues in any of their available conformers. These selected protein ensembles have at least five conformers in the database to assure a good estimation of the conformational variability ^57^. Only the pair of conformers showing the maximum Root Mean Square Deviation (RMSD) along all the ensemble was considered in this set. IDPs dataset consisted on 93 protein NMR ensembles selected from the PDB database. These ensembles have more than 40% of disordered positions as predicted by NMR-ESpritz ^58^ and Mobi 2.0 ^59^. Ordered set of proteins showed low levels of disorder as predicted with ESpritz X-ray (see Figure 3 and Figure S3).

### Structurally constrained substitution pattern estimation

In Figure S2 we resumed the workflow to analyse structurally constrained sites and physical contacts. For each conformer and each protein in both datasets (for the disordered dataset we considered all the NMR available conformers and for the ordered dataset we used those corresponding for the maximum RMSD according to CoDNaS), the SCPE model of protein evolution was run ^24^. SCPE derives site-specific substitution matrices using evolutionary simulations under neutral conditions for protein fold conservation ^46,60^. Briefly, it uses energetic calculations to evaluate the structural perturbation introduced by non-synonymous substitutions in the simulation process. Using maximum likelihood estimations (ML), it is possible to compare SCPE matrices with models lacking structural information such as JTT ^36^, Dayhoff ^61^ and WAG ^38^. Site-specific ML calculations were performed with the HYPHY package ^62^. The alignments used for the ML analysis were obtained from HSSP ^63^ database. Neighbour-joining distance phylogenetic trees were obtained with the Phylip ^64^ package. To define whether a site was structurally constrained (SC) Akaike information criteria (AIC) coefficient was used ^65^ and a ranking for the estimated models made using ΔAIC ^66^ in which models having ΔAIC <=2 have a substantial support, those where ΔAIC is between 4 and 7 have a intermediate support, while those with ΔAIC > 10 have no support. Tertiary contacts were estimated considering the distance between two non-contiguous residues having the van der Waals spheres of each residue side chain heavy atoms below 1.0 Å. Long-range inter-residues contacts were estimated using same definition but considering ±5 residues of a given residue.

## Acknowledgments

GP and MSF are CONICET researchers and JM and AMM are PhD and Postdoctoral fellows of the same institution.

This work was supported by Universidad Nacional de Quilmes (PUNQ 1004/11) (GP), Agencia de Ciencia y Tecnología (PICT-2014-3430) (GP) and COST Action (BM1405) NGP-net (SCET). This project has received funding from the European Union’s Horizon 2020 research and innovation programme under the Marie Sktodowska-Curie grant agreement No 778247 (IDPfun). The funders had no role in study design, data collection and analysis, decision to publish, or preparation of the manuscript.

